# Gradient of Tactile Properties in the Rat Whisker Pad

**DOI:** 10.1101/2020.02.27.967778

**Authors:** Erez Gugig, Hariom Sharma, Rony Azouz

## Abstract

The array of vibrissae on a rat’s face is the first stage in a high-resolution tactile sensing system. Progressing from rostral to caudal in any vibrissae row results in an increase in whisker length and thickness. This in turn may provide a systematic map of separate tactile channels governed by the mechanical properties of the whiskers. To examine whether this map is expressed in a location-dependent transformation of tactile signals into whisker vibrations and neuronal responses, we monitored whiskers’ movements across various surfaces and edges. We found a robust rostral-caudal gradient of tactile information transmission in which rostral shorter vibrissae displayed a higher sensitivity and a bigger differences in response to the different textures, whereas longer caudal vibrissae were less sensitive to the different textures. Nonetheless, we found that texture identity is not represented spatially across the whisker pad. Based on the responses of first-order sensory neurons, we found that neurons innervating rostral whiskers were better suited for textures discrimination, whereas neurons innervating caudal whiskers were more suited for edge detection. To examine the functional role of this organization, we monitored the whisking activity of awake rats foraging for food. We found a caudal-rostral gradient of whisking angle movements in which the longer caudal vibrissae spanned a larger space making them more suitable for object localization, whereas the rostral shorter vibrissae hardly moved. These results suggest that the whisker array in rodents forms a sensory structure in which different facets of tactile information are transmitted through location-dependent gradient of vibrissae on the rat’s face.

## Introduction

Facial whiskers, or vibrissae, are found in many mammalian species. They project outwards and forwards from the nose of the animal to form a tactile sensory array that surrounds the head [1]. This array is composed of two functionally distinct whisker systems - the long, moveable macrovibrissae which are actively swept across objects and surfaces in a rhythmic forward and backward motion called whisking [2-4], and shorter, non-moveable microvibrissae [5-7]. In rodents, the macrovibrissae form a two-dimensional grid of five rows on each side of the nose, where each row is composed of between five and nine whiskers [5], whereas the shorter microvibrissae have a higher density and are more anterior.

Rodents use their whiskers to detect and distinguish a variety of tactile features in their environment [8] including edge position [9, 10], shape [5, 11], aperture and gap width [12], and texture discrimination [7, 13-18]. Despite a long history of morphological, physiological and behavioral work indicating that the whiskers are functionally distinct (Hartmann’s [19, 20] and Moore’s [21, 22] groups), many neurophysiological studies assume that the whiskers are functionally equivalent, and that the array forms a diffuse spatial sensor. However, the evolutionarily conserved spatial arrangement of whisker arrays may have implications for probing the environment [5].

A few studies have addressed the functional architecture of the mystacial pad (i.e., the vibrissae pad). Several studies [5, 23] have suggested, based on behavioral data, that the mystacial macrovibrissae row is a “distance decoder”. Its presumed function is to derive head centered object shapes at the various angles represented by vibrissae rows while the microvibrissae are involved in object recognition tasks [7]. In recent years it became more accepted that the vibrissae in different regions of the array are not interchangeable sensors, but rather functionally grouped to acquire particular types of information about the environment [20]. Further support for this idea comes from a few studies showing a gradient in whiskers length and whisking movement range. Thus, the caudal whiskers are longer and move at larger angles than rostral whiskers [5, 13, 24-26]. An extension of this concept lies also in the “kinetic-signature” hypothesis [21, 27, 28], in which each whisker’s intrinsic dynamics (including but not limited to resonance) governs its texture-related response. According to this view, texture identity is represented spatially across the whisker pad. Only a specific set of textures will cause a group of whiskers to vibrate at their distinct natural frequency, making a set of whiskers selective for these particular textures, thus splitting the tactile vibration signals into labeled frequency lines in the cortex. These types of hypotheses are still a matter of debate in the field.

Given the changes in whisker properties along the mystacial arcs, we sought to examine the functional and behavioral implications of this gradient. The present study combines morphological, physiological and behavioral data to describe the relative functional contributions of individual whiskers in the mystacial pad. We suggest that the mystacial macrovibrissae form a gradient of tactile information transmission in which longer caudal vibrissae are mainly involved in active edge localization, whereas the rostral shorter vibrissae transmit both edge collision and texture coarseness information. This structure has functional significance for tactile exploration and navigation.

## MATERIALS AND METHODS

### Animals and Surgery

Two types of experiments were done in the current study: First, Acute anaesthetized rats. Second, Awake behaving rats. All experiments were conducted in accordance with international and institutional standards for the care and use of animals in research. For acute experiments, male Sprague Dawley rats (250-350 gm) were used. Surgical anesthesia was induced by urethane (1.5 gm/kg i.p.) and maintained at a constant level by monitoring forepaw withdrawal and corneal reflex. Extra doses (10% of the original dose) were administrated as necessary. Atropine methyl nitrate (0.3 mg/kg i.m.) was administered after general anesthesia to prevent respiratory complications. Body temperature was maintained near 37 °C using a servo-controlled heating blanket (Harvard, Holliston, MA). After placing the subjects in a stereotactic apparatus (TSE, Bad Homburg, Germany), an opening was made in the skull overlying the TG, and tungsten microelectrodes (2 MΩ, NanoBio Sensors, Israel) were lowered according to known stereotaxic coordinates of the TG (1.5-3 ML, 0.5-2.5 AP; [29, 30] until units drivable by whisker stimulations were encountered. The recorded signals were amplified (x1000), band-pass filtered (1 Hz-10 kHz), digitized (25 kHz) and stored for off-line spike sorting and analysis. The data were then separated into local field potentials (LFP; 1-150 Hz) and isolated single-unit activity (0.5-10 kHz). All neurons could be driven by manual stimulation of one of the whiskers, and all had single-whisker receptive fields. Spike extraction and sorting implemented MClust (by A.D. Redish available from http://redishlab.neuroscience.umn.edu/MClust/MClust.html), which is a Matlab-based (Mathworks, Natick, MA) spike-sorting software. The extracted and sorted spikes were stored at a 1 ms resolution, and PSTHs were computed.

### Whisker stimulation

We replayed whisker movements across different surfaces by covering the face of a rotating cylinder with several grades of sandpaper with different degrees of coarseness and rotating the wheel against the whiskers (Fig. 1A). The wheel face was placed so that the macrovibrissae rested upon it (Fig. 1A). The wheel was placed to mimic rostral-caudal whisker movement during head-movement. The head velocities associated with rat exploration were taken from Lottem and Azouz [18].The velocity was controlled using a DC motor driven at approximately 39 mm/sec to replicate median *head velocity*. The 30 mm diameter wheel was driven by a DC motor (Farnell, Leeds, UK). We employed surfaces of different grades (from coarse -grained to fine-grained (the numbers in the parentheses indicate the average grain diameter): P120 (125 µm), P150 (100 µm), P220 (68 µm), P400 (35 µm), P600 (26 µm), P800 (22 µm), P1200 (15 µm). These grades were chosen in accordance with previous studies [16, 31, 32]. For each texture, we recorded continuously for about 2 min. per texture of the rotating cylinder. To examine TG neuronal responses to edges, we attach a 5 mm edge to the face of the wheel. We divided the data into segments in which each segment contained whisker colliding with the edge and texture. Each revolution of the wheel lasted for about 3 sec. and no noticeable adaptation in neuronal firing rates were detected.

**Figure 1.**
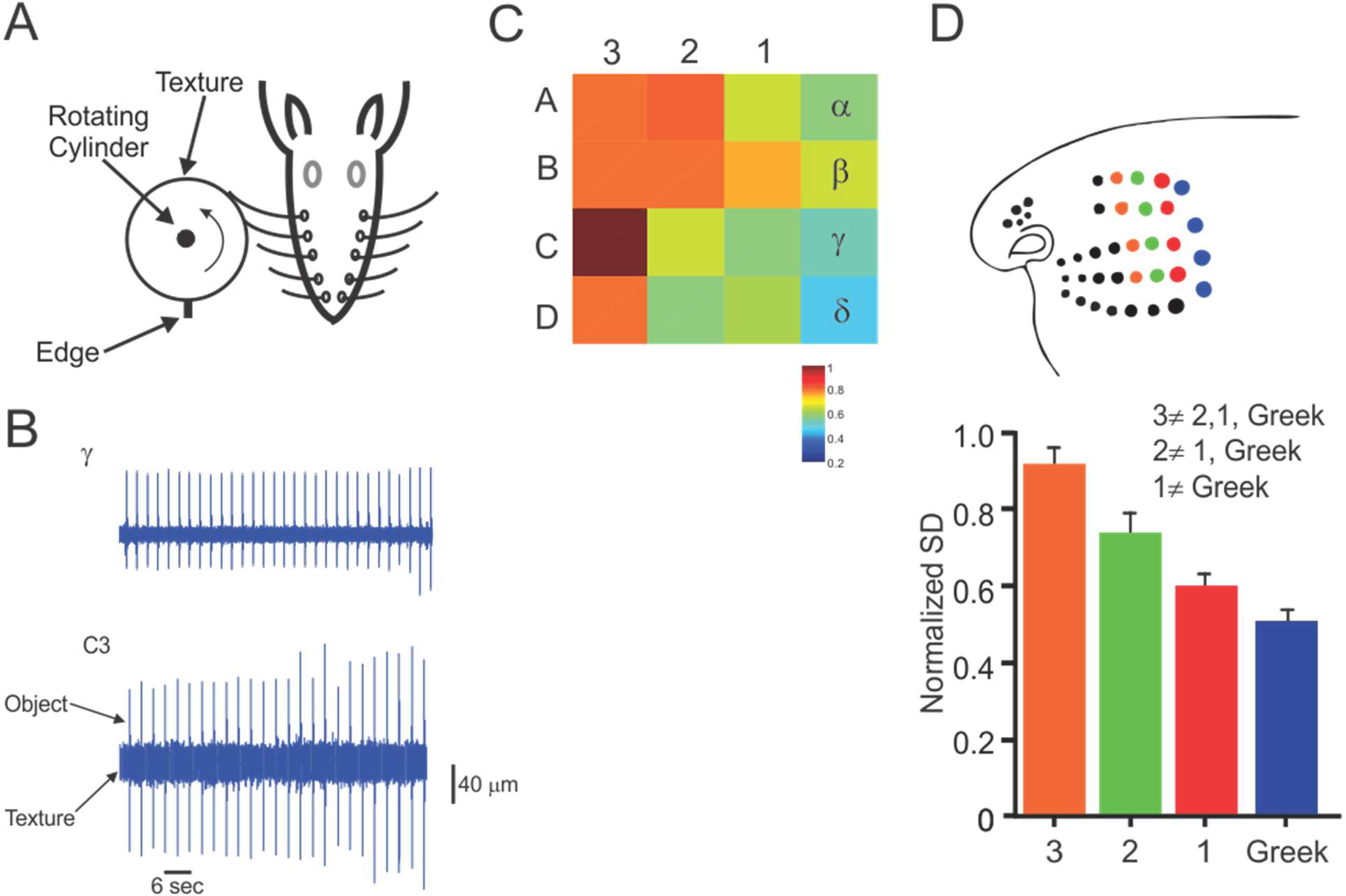
Differential mechanical characteristics of different whiskers arcs. (A). Experimental design. The whiskers are contacting a rotating cylinder covered with textured sandpaper and an edge. (B). Example of two outermost whiskers’ vibrations in response to texture (P220) and edge. (C). An heat map of response SD to all textures across all whiskers. Each pixel denotes the normalized whisker vibrations SD to the highest value in the map. (D). The upper panel shows the recorded Arcs. The lower panel shows the mean and SD of each of the arcs in C. The inequality sign indicates a statistically significant differences between the various arcs.

Whisker displacements transmitted to the receptors in the follicle were measured by an infrared photo-sensor (resolution: 1 µm; Panasonic: CNZ1120) placed about 2 mm from the pad. The voltage signals were digitized at 25 KHz and amplified (See Lottem and Azouz, 2008, for principles of sensor operation and a description of sensor calibration). In some of recordings (where mentioned), whisker images were acquired with Mikrotron CoaXPress 4CXP camera at 1600 frames/sec; 4 Megapixel resolution. The camera was installed above the arena and thus produced an overhead view. We rotated the texture-covered cylinders at velocities corresponding to head movements (see above).

### Behavioral paradigm and setup

To examine the functional role of the different macrovibrissae, we devised a behavioral task in which rats (male Sprague Dawley rats (250-350 gm) were used. *n* = 6) foraged for a food pellet on top of a round elevated platform (Fig. 8A; approximately 4 cm in diameter) in random locations at the edge of the arena. All animals were kept in a 12-h dark/light cycle at 22°C, 40-60 % humidity, with unrestricted access to water and food (unless otherwise noted); tested during the dark (active) part of their daily cycle; and handled prior to being placed in the experimental arena. The task was performed in the dark. Data collection took place in a 40 × 25 cm rectangular viewing arena with a glass floor and side walls. It was illuminated from below by an infrared light (15×15cm; Fig. 8A). Behaving rodents (*n* = 6) completed their task and picked up the food pellet. Whisker images were acquired at 200 frames/sec; 4 Megapixel resolution. The camera was installed above the arena and thus produced an overhead view. Due to the limitation of the camera, we were able to record only an area of 20×20 cm in proximity of the food pellet.

**Figure 2.**
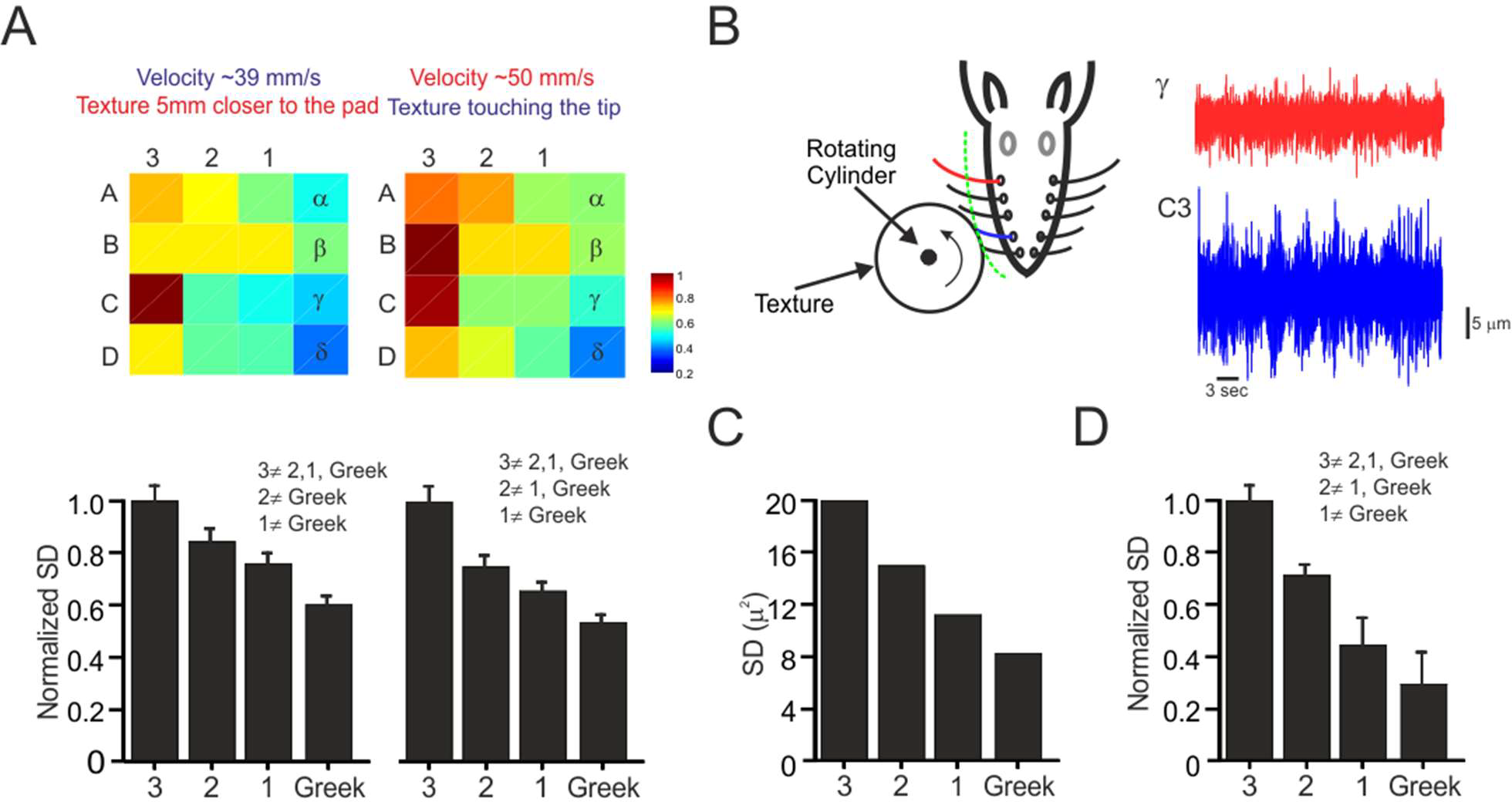
Robustness of whiskers’ properties map. (A). To examine to robustness of the map we changed the distance of the wheel to the pad (Texture 5mm closer to the pad; upper left panel), and wheel velocity (Velocity ∼50 mm/s; upper right panel). The lower panels show the mean and SD of the each arc in the upper panels. The inequality sign indicates a statistically significant differences between the various arcs. (B). Experimental design. The whiskers are contacting a rotating cylinder covered with textured sandpaper at the same distance. Example of two outermost whiskers’ (C3 and γ) vibrations in response to texture (P220). (C). The panel show the mean and SD of each of the arcs in B. (D). Normalized mean and SD of each of the arc in all animals (*n* = 5). The inequality sign indicates a statistically significant differences between the various arcs.

**Figure 3.**
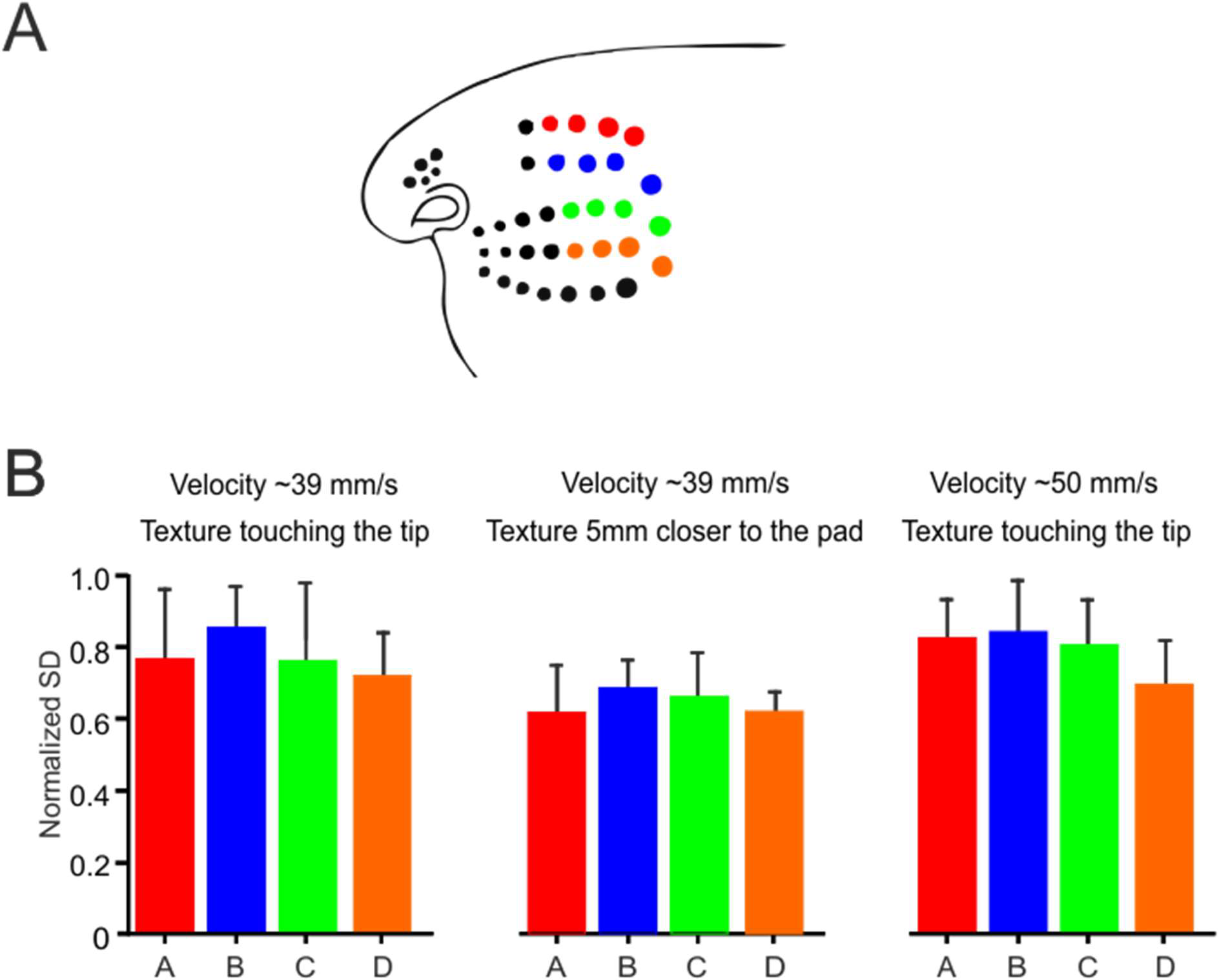
Mechanical characteristics of different whiskers rows are not different from each other. (A). The panel shows the recorded rows. (B). The panels show the mean and SD of each of rows in (A) at different distances from the pad and at different wheel velocity. The biomechanical properties of the whiskers across the different rows and in the different conditions are not significantly different from each other.

**Figure 4.**
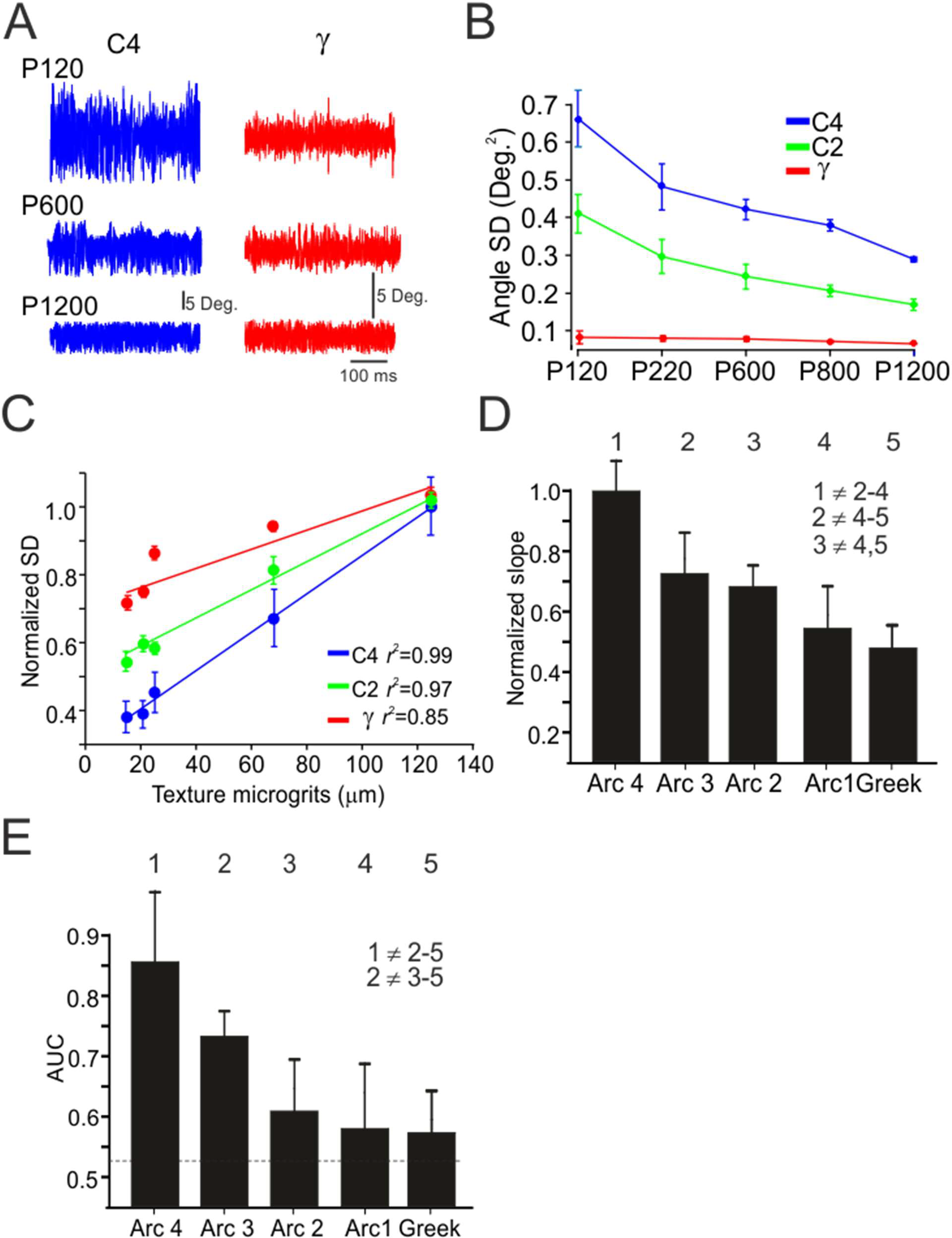
Response characterization of the different whisker rows to different textures. (A). Example of two outermost whiskers’ (C4, γ) vibrations in response to textures (P120, P600, P1200). (B). Quantification of 3 whiskers’ vibrations SD in response to different textures. (C). Linear regression fit of the normalized SD of whiskers vibrations in response to several textures (*n* = 5; each texture from B is represented by the mean particle size). (D). Normalized slopes of the linear regression fit for all whiskers. The numbers indicate the different arcs. The inequality sign indicates a statistically significant differences between the various arcs. (E). AUC for texture discrimination for the different arcs.

**Figure 5.**
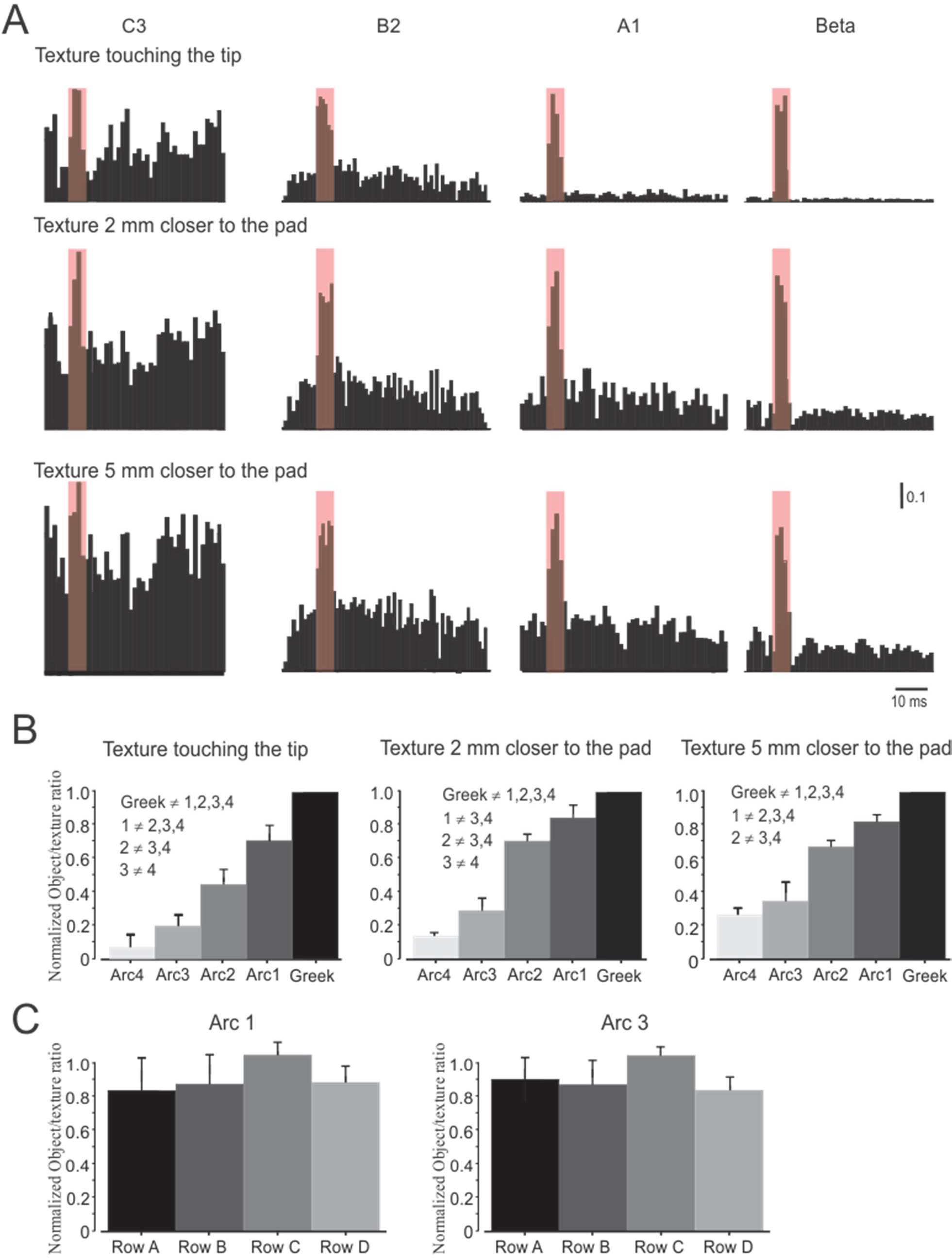
TG responses of different whiskers: (A). Typical examples of PSTHs from different arcs aligned to edge contact shown from caudal (right) to rostral (left). The red transparent square demarcates edge related responses. The upper, middle, and lower panels show the responses of the neurons at different distances of texture wheel from the pad. (B). Normalized firing rate ratio (the ratio between firing rates in response to edge and responses to textures normalized to the highest value across arcs) from the different arcs and at different conditions. (C). Normalized firing rate ratio (the ratio between firing rates in response to edge and responses to textures normalized to the highest value across rows) from two different rows. Scale bar of the PSTHs refers to firing probability at a given bin.

**Figure 6.**
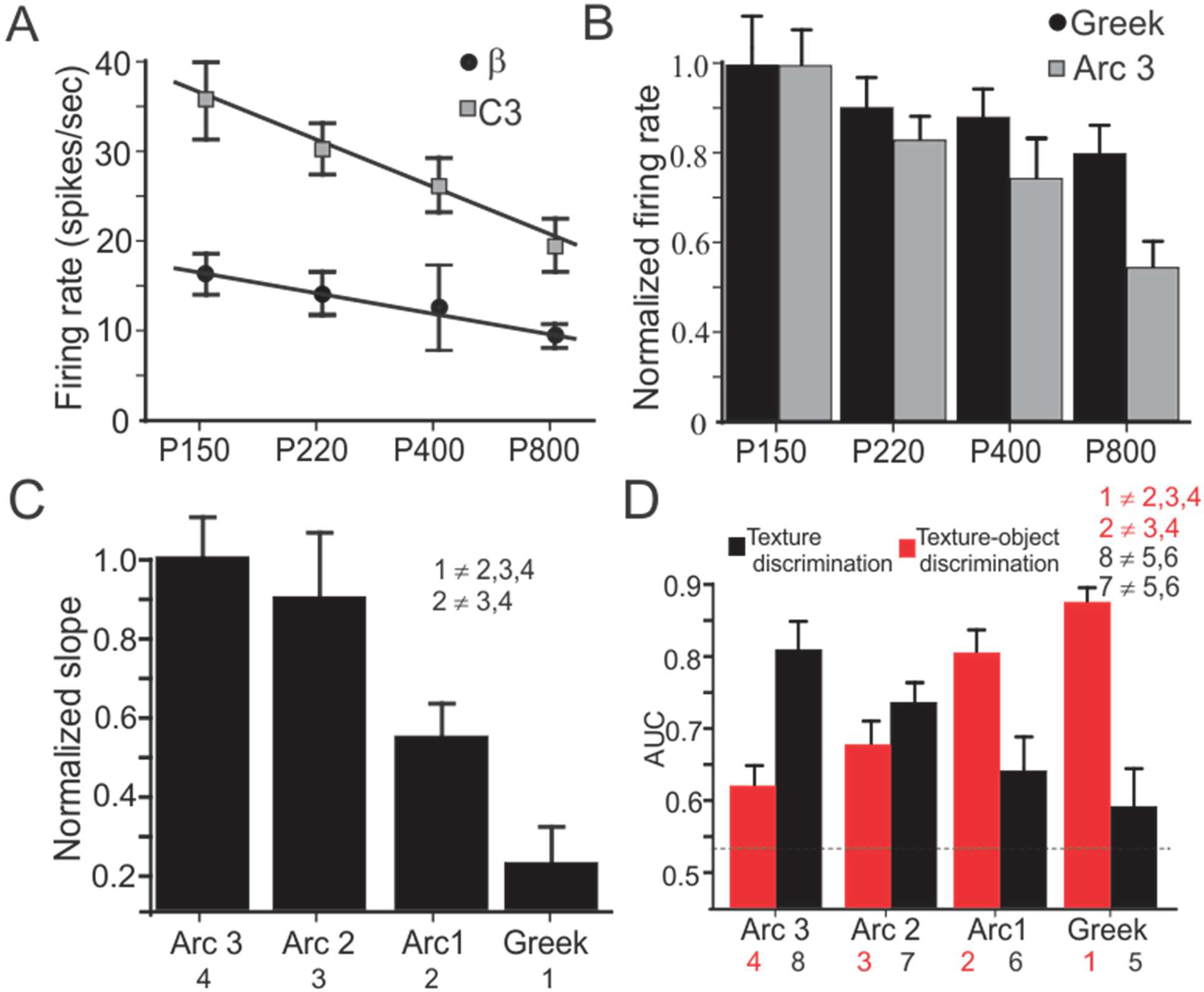
A gradient of textures and edge-texture discrimination capabilities. (A). The relationship between texture coarseness and TG firing rates in two arcs. (B). Normalized firing rates (normalized to the maximal firing rates) for all neurons innervating Greek and Arc 3 whiskers in response to the different textures. (C). Normalized slopes of the linear regression fit for normalized firing rates vs. texture microgrit (see Fig. 4C). (D). AUC for texture discrimination (black bars) and texture-edge discrimination (red bars) for the different arcs. The numbers indicate the different arcs. The inequality sign indicates a statistically significant differences between the various arcs.

**Figure 7.**
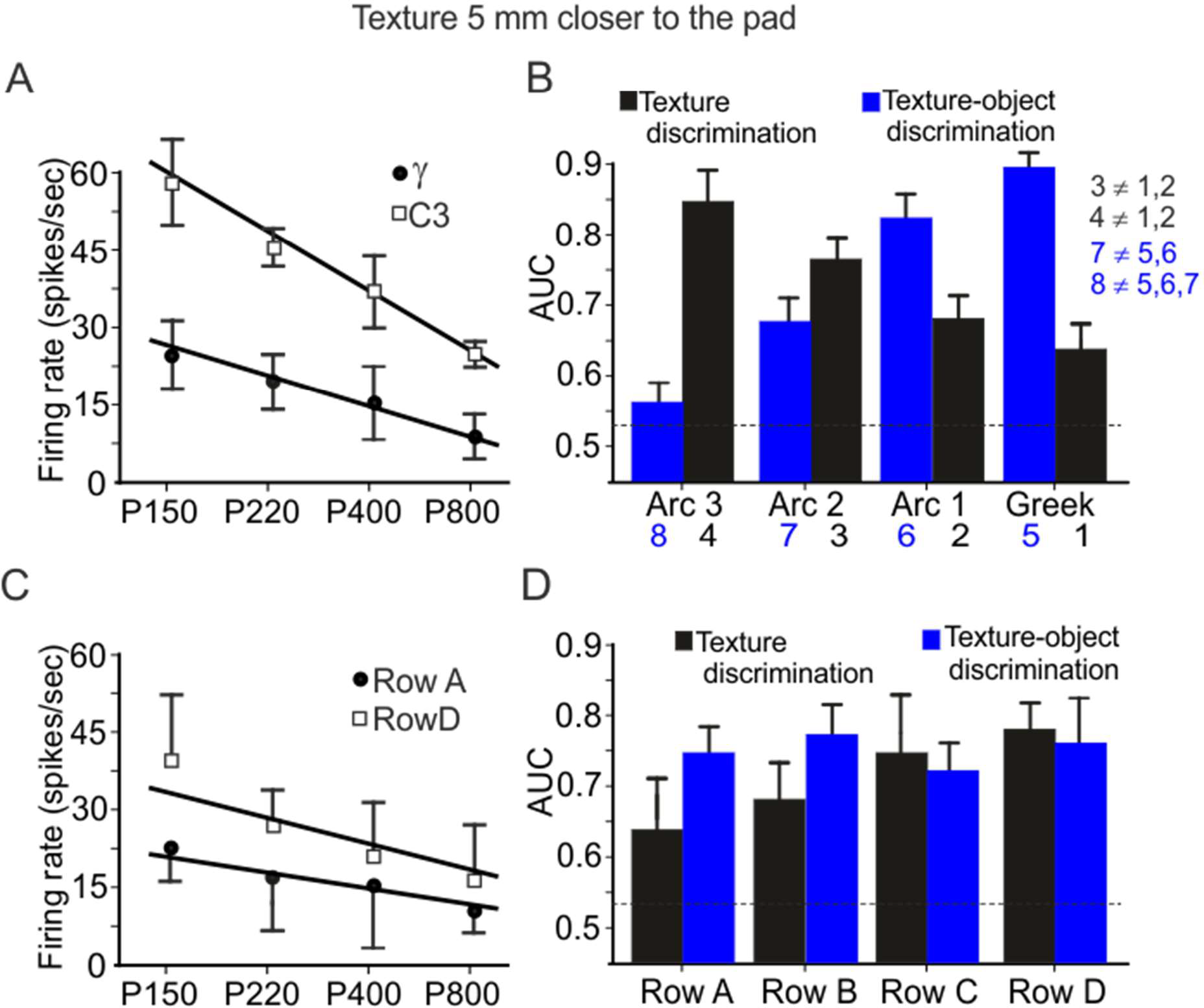
A robust gradient of textures and edge-texture discrimination capabilities in TG neurons. (A). Similar to Fig. 6A but textures are 5 mm closer to the pad. (B). AUC for texture discrimination (black bars) and texture-edge discrimination (blue bars) for the different arcs. (C). Similar to Fig. 6A but for rows. (D). Similar to B but for rows. The numbers indicate the different arcs. The inequality sign indicates a statistically significant differences between the various arcs.

**Figure 8.**
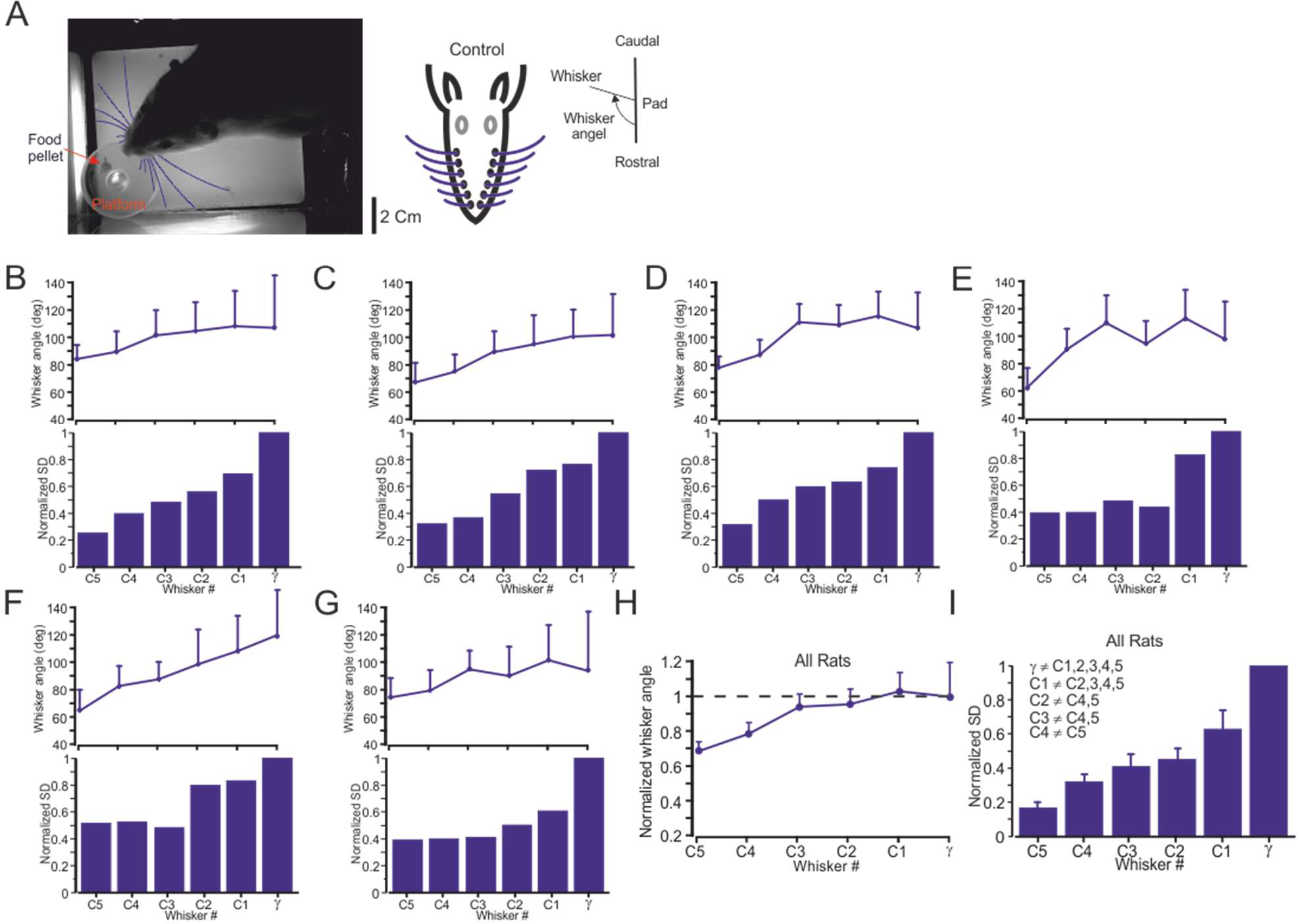
Differential whisking strategies during exploration. (A). Experimental design. Rats forage freely in the dark for a food pellet on a round elevated platform (see methods section). The whiskers are marked in blue. The right panel show the way whisker angle was calculated. (B-G upper panels). Mean and whisking amplitude for each whisker. The whiskers angle range is denoted by the error bars, which are indicated by the SD. (B-G lower panels). Whisking amplitude for all animals as indicated by the SD (normalized in each rat by whisker’s maximal angle/ γ). The caudal whiskers show a larger whisking amplitude. (H). Mean whiskers angle and amplitude for each whisker (*n* = 6) for all animals, normalized to the mean and SD of γ whiskers. (I). Whisking amplitude for all animals normalized to each animal γ whisker angle SD values. The # sign indicates statistically significant different groups.

The experiment was comprised of two stages: Training and handling, experimental sessions. During the Training and Handling phase (∼7 days) animals gradually learned to search for food pellets on the platform in the arena upon its opening. The animals were food deprived approximately 16-20 hours prior to training or experimental sessions in order to maximize food foraging during sessions. Animals began the experimental stages once they did not show stress symptoms (such as “freezing” in one of the corners) upon entering the arena and engaged rapidly the foraging task. Opportunistic recordings, each 20–40 sec in length, were taken of awake, unrestrained animals engaged in active, exploratory behavior. *For better visualization, all whiskers rows were cut, except row C*. Each recording was initiated by an experimenter viewing the camera scene on a monitor window and pressing a trigger when the animal entered the field of view. We monitored whiskers movement while the animals approached the food pellet on top of the platform. We monitored whisker movements, using high speed camera, during the approach to the food pellet on top of an elevated platform. All the movies were analyzed using the Janelia whisker tracker software [33]. Recording stopped when the camera memory was full, the animal stopped exploring, or became stressed. The top-down view clips from each rat searching throughout the arena for the food pellet on top of the platform were selected for analysis, or portions thereof, using the following selection criteria developed in Grant et al. [6]: (1) the rat head was clearly in frame; (2) both sides of the face were visible; (3) the head was level with the floor; (4) the whiskers were not in contact with a wall.

### Data analysis

To examine the influence of whisker identity on responses to edges and textures (Fig. 1A,B), we separated edge and texture related stimulation epochs by identifying big excursion (mean±3SD) in whisker vibrations during wheel rotation. We then aligned the whisker responses and the corresponding neuronal responses to create PSTHs (Fig, 5A). Once PSTHs were created, we defined manually the temporal margins of edge responses (Fig. 5A). To determine the ratios between edge and texture responses, we calculated the corresponding firing rates.

The significance of the differences between measured parameters was evaluated using one-way analysis of variance (ANOVA). When significant differences were indicated in the F ratio test (*p*<0.05), the Tukey method for multiple comparisons was used to determine those pairs of measured parameters that differed significantly from each other within a group of parameters (*P*<0.05 or *P*<0.01). The results are presented as the mean ± standard deviation of mean (SD). Error bars in all the figures indicate the SD unless otherwise noted. To avoid cluttering in some of the graphs we use single-sided error bars.

### Receiver operating characteristics analysis

We used signal detection theory (receiver operating characteristics, ROC analysis, [34], to compute the probability that an ideal observer could accurately determine the differences among the different textures based on neuronal activity. For each measured texture pair, an ROC curve was constructed. The ROC curve is a two-dimensional plot of hit probability on the ordinate against false-alarm probability on the abscissa. To transform raw data into a measure of discriminability, we analyzed the distributions of neuronal firing rates *<u>across trials</u>*. Green and Swets (1966) showed that the area under the ROC curve (AUC) corresponds to the performance expected of an ideal observer in a two-alternative, forced-choice paradigm, such as the one used in the present analysis. The ROC curve was calculated for the firing rate of a single neuron as a function of texture. We then averaged all AUC values of all neurons, all texture pairs in the different rows or arcs.

The firing rate in trial *k* is the spike count 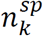 in an interval of duration *T* divided by *T*

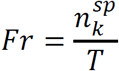

The length *T* for texture signal was set to *T*=50 ms and for edges *T*=10 ms.

To measure the significance level of P(correct) in the ensemble of TG neurons, we took all possible textures comparisons for all neurons and shuffled the trials across the different stimuli. We then repeated this procedure 500 times. The significance level was set at 90% of this population, namely 0.53 (Fig. 4E, 6D, 7B,D, dashed lines).

## Results

We examined the differential role of the macrovibrissae in the transmission of tactile information, and quantitatively evaluated the mechanical and neuronal mechanisms underlying this process. To begin with, we sought to examine whether the mystacial pad has a location-dependent differentiation of transmission of “what” information. We took several approaches to address this issue. *First*, we determined a gradient of whiskers’ biomechanical characteristics across the pad. *Second*, we examined whether this gradient is expressed in neuronal responses to textures. To evaluate whether the different biomechanical properties of the whiskers play a role in tactile information transmission, we replayed passive whisker movements across different surfaces by covering the face of a rotating cylinder with several grades of sandpaper with different degrees of coarseness. The cylinder face was placed orthogonally to the vibrissae so that the vibrissa rested upon it (Fig. 1A). These surfaces were placed at two different distances from the pad (Whisker tip and 5 mm closer to the pad) and at two different wheel velocities (39 mm/sec and 50 mm/sec). Vibrissae movements were measured in response to sandpapers having two different grades: P220, P400. However, since we got similar results for all textures (not shown), we combined all these results together.

We examined the influence of the biomechanical properties of the vibrissae on the transformation of surface coarseness into whisker vibrations by measuring across several anaesthetized animals (n = 5) for each vibrissa. In Fig. 1B, we show an example of whisker vibrations in two vibrissae in response to P220 texture and an embedded edge. These panels show that rostral shorter, slender vibrissae expressed increased range of vibrations for textures and edges, whereas the caudal vibrissae presented lesser variance. To establish a functional map of the mechanical properties of the vibrissae we first calculated the SD of each of the measured whisker vibrations in response to all studied textures. We then, normalized each texture SD to the maximal value across all whiskers. We then averaged the normalized values for each whisker in the map. That is, the pixels values in the maps in Fig. 1C are averages of normalized SD for all textures across all whiskers. A relatively smaller variance indicates a larger attenuation of the texture signal. This analysis revealed a location-dependent gradient in which whisker vibrations were transmitted more robustly through rostral whiskers (Fig. 1C). Each pixel in this plot represents an average of numerous vibrissae [Greek:(α: *n* = 5; β: *n* = 5; γ: *n* = 3; δ: *n* = 5); arc 1: (A: *n* = 5; B: *n* = 5; C: *n* = 5; D: *n* = 5); arc 2: (A: *n* = 5; B: *n* = 5; C: *n* = 5; D: *n* = 5); arc 3: (A: *n* = 5; B: *n* = 5; C: *n* = 5; D: *n* = 5); Due to size limitation of the sensor and the short length of rostral whiskers, we did not record whiskers’ displacements and corresponding neuronal responses to whiskers rostral to arc 3]. We then collapsed all values in each arc across all animals and examined the statistical validity of the R-C gradient (Fig. 1D upper panel). We found statistically significant differences between most arcs (Fig. 1D lower panel). These results suggest a caudal-rostral functional gradient that arise from the biomechanical differences of the sensing organs (i.e., the whiskers).

To examine the robustness of this gradient, we took several approaches: *First*, we changed wheel velocity and the proximity of the wheel to the pad. We found that decreasing the distance between moving textures and the pad resulted in a modification of the map, while keeping the rostral-caudal gradient nearly constant (Fig. 2A, left panel panel). Similarly, when increasing texture velocity, we saw minor changes in the map; nevertheless there were no major changes in the rostral-caudal gradient (Fig. 2A, right panel). In both of these conditions, we collapsed all values in each arc across all animals and found statistically significant differences between most arcs (Fig. 2A lower panels). *Second*, when a rat is whisking against a textured surface, due to whiskers having different lengths, the stimulation might occur at different locations along the shaft for different whiskers. Therefore, we changed the proximity of the wheel to the pad to keep the distance from the pad constant. Figure 2B depicts the experimental paradigm and whiskers C3 and γ vibrations in response to P220 at the same distance from the pad. We then quantified the SD of whisker vibration in this animal and found a gradual decrease of the whisker vibrations as a function of whisker identity (Fig. 2C). Averaging across all animals from both sides of the head (*n* = 3) we found a consistent decrease in whisker vibrations for more caudal whiskers (Fig. 2D). Finally, to examine whether this gradient exists in the dorsal-ventral plane, we repeated the same analysis by averaging across rows. Figure 3 shows that the whiskers along the rows are not significantly different in their responses to textures.

To examine whether the gradient in whiskers’ mechanical properties is instrumental in texture discrimination we compared whisker vibrations to five textures (P120; P220; P600; P800, P1200) in the different arcs. Examples of whisker vibrations in C4 and γ whiskers is shown in Fig. 4A. The panel show that the range of whisker vibrations is higher for all textures in C4 whisker vs. γ whisker. Verification of this notion is shown in the average for C4, C2 and γ whiskers (*n* = 5) in Fig. 4B. We quantified the range of whisker vibration in response to the different textures by calculating the SD. The figure shows that the rostral whisker respond with a higher range of whisker motion than caudal whisker. Moreover, the difference in SD between the textures is larger in the rostral whiskers. Finally, there exists a gradual relation between texture coarseness and whisker variance across the whisker pad i.e. coarser and finer surfaces are expressed in a higher and lower SD, respectively. This implies for all whiskers. To quantify the difference between textures in the different whiskers, we calculated the linear regression fit for the normalized SD for each whisker as a function of mean particle diameter in each texture (Fig. 4C). We found that the slope was steeper in C4 than in γ whiskers. Figure 4D. shows this in all recorded whiskers. The normalized slopes show that rostral whiskers display a bigger difference between textures than caudal whiskers. Finally, we employed ideal observer analysis to quantify the discriminative power of each of the whiskers. We divided the continuous whisker signals into multiple segments (500 ms in duration) and used the AUC measure and calculated for each whisker, the average AUC across all texture combinations. We found that rostral whiskers can better discriminate between the different textures (Fig. 4E). We conclude that the mechanical properties of the pad whiskers enables them to transmit differential aspects of tactile information.

We sought to test how this gradient directly translates into neuronal activity in first-order sensory neurons by examining the relationship between neuronal responses to textures and edges in the different whiskers. We took several approaches to examine this gradient: *First*, we recorded responses to step whisker deflections from 71 TG neurons obtained from 16 adult rats. As previously described [30, 35-38], such neurons respond to step stimuli with either a phasic response, firing only to stimulus onset/offset, or a phasic-tonic response, firing both at stimulus onset/offset and throughout the stimulus hold period. These firing patterns are conventionally used to characterize neurons as either rapidly adapting (RA) or slowly adapting (SA), respectively. We did not find any significant bias in neuronal types towards any of the arcs (Table 1). *Second*, we recorded extracellularly from 57 trigeminal ganglion (TG) neurons (Greek: *n* = 13; arc 1: *n* = 15; arc 2: *n* = 9; arc 3: *n* = 10; arc 4: *n* = 11). The experimental paradigm is shown in Fig. 1A. An example of neuronal responses of four separate TG neurons to a rotating wheel covered with textures and an edge is shown in the upper panel in Fig. 5A. We initially, divided the continuous neuronal recordings (Fig. 1B) into segments which contained an epoch preceding response to edges, responses to edge and an epoch containing responses to textures (Fig 5A, red boxes). Delineation of these epochs within each segment was done using the criterions from whisker signals (see methods section). Once these segments were created, we created PSTHs for each neuron for the different conditions (Fig. 5A). The PSTHs of the neurons indicated that the caudal whiskers’ neurons responded mainly to edges while progressively more rostral whiskers’ neurons responded to both edges and texture stimuli. This was further verified by calculating the normalized ratio between discharge probability in response to edges and textures (normalization was done as the ratio between each whiskers value and the maximal value, which was always the Greek arcs). Figure 5B left panel shows that this ratio decreased gradually from caudal to rostral, since texture-associated firing increased gradually when moving to whiskers that are more rostral. To examine the robustness of this phenomenon, we repeated these experiments at several distances from the pad (Figure 5A, middle and lower panels). Decreasing the distance from the pad resulted in an increase in neuronal firing probability. Nonetheless, the ratio decreased gradually from caudal to rostral whiskers (Figure 5B, middle and right panels). Finally, we compared the firing rates across the different *rows* and didn’t find any statistically significant differences in their firing rates (see methods section for statistical significance of the differences between measured parameters; Fig. 5C). Our results indicate that irrespective of surface distance, the ratio between discharge rates in response to edges and textures does not change qualitatively (Figure 5B, middle and lower panels).

**Table 1.** Distribution of neuronal types across all whiskers.

| Neuronal type \ Arc | Arc |  |  |  |  |
| --- | --- | --- | --- | --- | --- |
| | Greek ($n = 20$) | 1 ($n = 15$) | 2 ($n = 16$) | 3 ($n = 15$) | 4 ($n = 5$) |
| RA | 9 | 8 | 9 | 7 | 3 |
| SA | 11 | 7 | 7 | 8 | 2 |

To examine the influence of whiskers’ biomechanical properties on tactile information transmission, and whether these characteristics affect neuronal ability to discriminate between different textures as well as between textures and edges, we used the ideal observer approach (see methods) and calculated the AUC, which is a measure of discriminability between two distributions, for the firing rates for each of pair of textures and edges in the different whiskers for all neurons. We initially divided the data into two epochs: neuronal responses to fine grained textures and edges (Fig. 5A). We then calculated the firing rates for each texture and edge epoch. An average firing rates (for each texture epoch in the neuronal response) of two neurons taken from the same animal for β and C3 whiskers is shown in Fig. 6A. The panel show a steeper change in firing rates across textures in C3 neurons. Whereas, β neurons changed only slightly their firing rates to the different textures. We then, calculated the normalized firing rates (normalized to highest firing rates across all whiskers) and found a steeper slope for firing rates in Arc3 vs. Greek (*n* = 23). To quantify the relations between surface coarseness and firing rates in all neurons across all arcs, we calculated the slope of the linear regression between grit size (P150 = 100 µm; P220 = 68 µm; P400 = 35 µm; P800 = 22 µm) and normalized firing rates for each neuron. We then, normalized the slope of each neuron to the largest slope in the sample (*n* = 57), and divided the values to the corresponding arcs. We found a steeper slope for the rostral whisker then the caudal whisker, suggesting that neurons innervating rostral whiskers are better suited for discriminating between textures. To examine this premise we used the ideal observer approach and compared the mean AUCs across all textures pairs for all whiskers (*n* = 57). We found that TG neurons in rostral whiskers have significantly higher AUCs suggesting they perform better in fine grained texture discrimination (Fig. 6D, black bars). In contrast, comparing the firing rates of fine grained textures and edges revealed that TG neurons in caudal whiskers have significantly higher AUCs for texture-edge discrimination (Fig. 6D, red bars).

To examine the robustness of this phenomenon, we repeated these measurements in a different set of whiskers at a closer distance of the wheel to the pad. While we observed an increase in the firing rates (Fig. 7A), the gradient in texture and edge-texture discrimination persisted. Together, these results indicate a differential role for caudal and rostral whiskers in texture and edge discrimination. Specifically, the longer, more movable caudal whiskers are more suitable for distinguishing between collision with an edge and vibrations associated with texture and the shorter, less movable rostral textures are more suitable to discriminate fine surface details. Finally, we examined the role of the different *rows* in texture and texture-edge discrimination. Figure 7C shows that A3 neuron has a steeper dependence of firing rates on texture coarseness than D3 neuron. However, averaging across all neurons revealed that neurons in the different rows do not differ in their texture discrimination and edge-texture discrimination (Fig. 3F).

To examine the functional role of the whiskers in different arcs, we devised a behavioral task in which rats foraged for a food pellet in random locations at the edge of the arena illuminated with infrared light (Fig. 8A; 40 × 25 cm). The food pellet was located on top of a round platform (approximately 4 cm in diameter). The animals (*n* = 6; for better visualization, all whiskers rows were cut, except row C) searched for food in random locations. Each trial lasted 20-40 sec, which yielded in 4000-8000 frames. We monitored whisker movements, using high speed camera, during the approach to the food pellet on top of the platform.

Inspection of whisker position in the animal in Fig. 8A reveal that most rostral whiskers are directed forward to sense the incoming surroundings as the rat moved forward, whereas the caudal whiskers point more backward. This is shown in Fig. 8B-G for all 6 animals. To quantify whiskers’ movement range in the different arcs, we averaged all whisker angles in all frames and trials and calculated the mean and SD of the mean whiskers’ angle in each animal. Fig. 8B-G corroborates the notion that rostral whiskers are directed forward, whereas caudal whiskers point more backward. To quantify the angle range in each whisker, we normalized each arc’s SD (Fig 8B-G, error bars) to γ’s SD. The lower panels show that the whiskers’ angle range is larger in longer posterior whiskers and decreases in shorter rostral whiskers. We performed these analysis on all six animals by normalizing each whisker’s position and range to that of γ’s. This normalization was performed on each frame in all trials. We found similar results for both average whisker angle and range across the different arcs (Fig. 8H). Using ANOVA (see methods section), we found significant decrease in whisking amplitude for progressively rostral whiskers (Fig. 8E). These results suggest location-dependent whisking ranges in rats foraging for food.

## Discussion

Rodents are strongly tactile animals who scan the environment with their macrovibrissae, and orient to explore edges further with their shorter and more densely packed, microvibrissae [5, 7, 26]. The facial vibrissae are organized into a pattern of several arcs and rows [5, 39]. We posit that this pattern could serve a functional role in the way animals sense the environment. Here, we focused on the function of this orderly array of facial whiskers and addressed the issue of the relationship between the structural features and functional operations of the vibrissae apparatus. We suggest that the mystacial whiskers form a gradient of tactile information transmission. In this scheme, longer caudal whiskers transmit mainly “where” information, whereas rostral shorter vibrissae transmit both “where” and “what” information. Our results suggest that whisker array in rodents forms a sensory structure in which different aspects of tactile information are transmitted through a location-dependent gradient of vibrissae on the rat’s face.

One major feature that stem from our results is that within a single whisk, rats may sample a region of an object at multiple spatiotemporal scales simultaneously. This may enable rodents to extract complex object features in a single whisk. The functional architecture of the pad ensures that vibrissae in different columns sample objects at different resolutions. Specifically, the rostral vibrissae sample objects with a higher spatiotemporal resolution than caudal vibrissae. This is consistent with the possibility that the rostral whiskers immediately surrounding the snout may serve as a high-acuity “fovea” during tactual exploratory behavior [2, 7, 40]. This hypothesis supports the notion whisking strategies are dependent on the behavioral task as well as the structure of pad. The morphology of the vibrissal array has critical implication on the nature of the neural computations that can be associated with extraction of edge location and features. Thus, the differential whiskers’ intrinsic properties in which length is critical suggests the possibility for a parallel transduction of edges’ location, shape and textures through independent sensory channels [41].

### Edge localization

Our results point to the fact that several factors such as whisker properties, motor commands, and neuronal response profiles all serve to influence behavioral strategies. We found an anterior-posterior gradient of whisking amplitude and lateral movement across the pad. Specifically, the longer caudal vibrissae span a larger space, whereas the rostral shorter vibrissae hardly move. Further support for these findings comes from the simple behavioral paradigm we used in the current study. We show that during free exploration for food pellets, the caudal whiskers move at a larger whisking amplitude, whereas the rostral whiskers hardly move (Fig. 8), suggesting that the caudal whiskers might play bigger role in active edge localization. It would appear that by the interplay between whisking amplitude and speed, rats could control the space and the speed by which it senses the environment. These strategies could also be understood as an active touch strategy aimed at prolonging contact and thereby aiding the extraction of information about surface characteristics such as texture [26]. Our results are consistent with the current literature suggesting that a significant component of whisking behavior is the “spread,” between the whiskers. Observed spread is shown to vary over the whisk cycle and to substantially decrease during exploration of an unexpected surface [26, 42, 43]. Because of larger whisking amplitude of the caudal whisker they will be the most critical in modulating whisker spread. The robustness of these results in the two behavioral conditions, suggest that these differential whisking characteristics may serve a functional role is tactile information transmission.

### Mystacial pad anatomy

The anatomical basis for the gradient in whisking may depend on the organization of the muscles in the mystacial pad in relation to the whisker follicles and surrounding tissue. Movement of the whiskers is controlled by the facial motor nerve, which innervates two classes of muscles: the intrinsic and extrinsic muscles. In awake animals, whisker retraction probably involves the activation of extrinsic facial muscles, while protraction involves the intrinsic muscles [2]. The differential whisking amplitude across the pad could result from the anatomical structure of pad muscles. That is, the superficial extrinsic facial muscles could pull more caudal whiskers further back during retraction, and having less of an impact on the more rostral whiskers. This will result in a larger movements of the more caudal whiskers. Alternatively, Haidarliu et al. [44] studied the nerve subdivision and the numerous mystacial pad muscles in the rat extensively and found a gradient in which the caudal whiskers are surrounded by larger intrinsic muscles and a larger subdivision of the extrinsic muscles. All these make the caudal vibrissae better suited for active edge localization.

In addition to this global whisking control, several studies have shown that the rat does have sufficient degrees of freedom of whisker control to effect some differential movement of either individual whiskers or whisker columns [2, 45-47]. This individual follicle control may be the basis for complex kinematics and rich variability in whisking behavior [13, 26, 42, 47-56].

### Texture discrimination

We found that whiskers, which are the first stage of tactile information translation, form a location-dependent gradient in which rostral whiskers show smaller attenuation in the transformation from texture coarseness to whisker vibrations, whereas caudal whiskers mainly transmit larger signals such as edge collision (Fig. 1-4). Further support for these finding comes from our lab [18] showing that the spectral contents of whisker vibrations in response to textures varies across arcs. This study show that rostral whiskers transmit more power in the high-frequency range, thereby enabling the transmission of stick-and-slip event more robustly. This further supports the notion that rostral whiskers are better suited for texture discrimination. On the other hand, caudal whiskers are better suited for transmitting lower frequencies; i.e., movement of the whiskers brushing against walls and edges and large deflection such as active edge contact. Subsequently, the whiskers’ biomechanical characteristics were manifested in the differential sensitivity of first-order sensory neurons.

We found an anterior-posterior gradient of tactile information transmission in which TG neurons of longer caudal vibrissae mainly transmit edge collision information, whereas TG neurons of rostral shorter vibrissae transmit both edge collision and texture coarseness information (Fig. 5-7). Our results suggest that these differential neuronal responses may stem from the mechanical properties of the whiskers and not from a change the neuronal characteristics of the mechanoreceptors of the different whiskers. Moreover, based on neuronal firing rates, we found that rostral whiskers’ neurons are better suited for finer details of the tactile environment, whereas caudal whiskers’ neurons are better suited for coarser details such as edges collision (Fig. 5-7). Several important implications arise from these results. Since all tactile information available to the whisker somatosensory system originates in these mechanoreceptors, it is conceivable that the gradient of tactile information transmission on the vibrissa pad is expressed as a somatotopic cortical map that describes multiple spatiotemporal scales extending along the representations of arcs of vibrissae, much like the resonant theory [22]. Moreover, we posit that being exposed continuously to these diverse inputs may shape differentially both the wiring and the properties of the constituent neurons all along the pathway up to the cortex [57].

While we found a gradient of mechanical properties along he whisker pad which were reflected as texture-related response, our results do not show that texture identity is represented spatially across the whisker pad. That is, we could not support the hypothesis that only a specific set of textures will cause a group of whiskers to vibrate at their distinct natural frequency, making a set of whiskers selective for these particular textures, thus splitting the tactile vibration signals into labeled frequency lines in the cortex.

### Methodological considerations

In the current study, whiskers were stimulated passively, thus our results may be accounted for by tactile inputs caused by head movements [18]. Nevertheless, the carriers of the tactile signal (stick and slip events) were preserved in both conditions [18, 58]. More importantly, the present work addressed a rather simple configuration in which the whiskers were passively dragged along a surface near-parallel to the mystacial pad, while exploration generally involves active whisking onto an edge located in front of the animal’s snout. Such configurations involve complex 3D whisker dynamics[53].

The whisking space covered by the whiskers during different behavioral modes is not confined to only to the R-C plane but whiskers’ motion is characterized by translational movements and three rotary components: azimuth, elevation, and torsion [48, 53]. However, it has been shown that during a protraction, while roll and elevation changes have strong effects on the trajectories of the vibrissal tips, the net effect on the volumetric search space of the rat is small (about 4%) [19]. In the current study, using one video camera which was place above the animals, we were able to analyze only the R-C plane. However, we believe that this is the major plane, rats sense their environment. Further support for this notion comes from our findings that no significant changes in biomechanical and neuronal response properties were observed in dorsal-ventral plane (see below).

Although we sought to determine the maximal range of each vibrissa movement, we did not examine systematically whisking kinematics, which has been studied thoroughly and is known to vary considerably depending on the behavioral task [26, 51, 59, 60]. Previous studies have found that sometimes the rostral whiskers move through larger angles than the caudal whiskers, while other times caudal whiskers move through larger angles than the rostral whiskers [19]. Therefore, our findings are only relevant for the behavioral paradigms used here, freely moving animals foraging for food.

## Acknowledgments

This work was supported by a grant from the Israel Science Foundation to RA and partially supported by the Helmsley Charitable Trust through the Agricultural, Biological and Cognitive Robotics Initiative of Ben-Gurion University of the Negev.

